# Convergence of modality invariance and attention selectivity in the cortical semantic circuit

**DOI:** 10.1101/2020.06.19.160960

**Authors:** Tomoya Nakai, Hiroto Q. Yamaguchi, Shinji Nishimoto

## Abstract

The human linguistic system is characterized by modality invariance and attention selectivity. Previous studies have examined these properties independently and reported perisylvian region involvement for both; however, their relationship and the linguistic information they harbor remain unknown. Participants were assessed by functional MRI, while spoken narratives (auditory) and written texts (visual) were presented, either separately or simultaneously. Participants were asked to attend to one stimulus when both were presented. We extracted phonemic and semantic features from these auditory and visual modalities, to train multiple, voxel-wise encoding models. Cross-modal examinations of the trained models revealed that perisylvian regions were associated with modality-invariant semantic representations. Attentional modulation was quantified by examining the modeling performance for attended and unattended conditions. We have determined that perisylvian regions exhibited attention selectivity. Both modality invariance and attention selectivity are both prominent in models that use semantic but not phonemic features. Modality invariance was significantly correlated with attention selectivity in some brain regions; however, we also identified cortical regions associated with only modality invariance or only attention selectivity. Thus, paying selective attention to a specific sensory input modality may regulate the semantic information that is partly processed in brain networks that are shared across modalities.

---

Modality invariance and attention selectivity are two properties that are characteristic of language communication. We understand linguistic contents, regardless of their presentation in text or speech (modality invariance). When we are exposed to different linguistic stimuli simultaneously, however, attending to auditory information often prevents the understanding of information presented visually (attention selectivity).

Previous studies have reported modality-invariant brain activity, associated with single-word processing (Booth et al., 2002; Marinkovic et al., 2003), sentence comprehension (Carpentier et al., 2001; Jobard et al., 2007), and story comprehension (Deniz et al., 2019; Regev et al., 2013) In particular, Deniz et al. (2019) quantitatively estimated common semantic information across visual and auditory modalities, even after excluding the effects of other linguistic and sensory features. Modality-invariant linguistic information is likely represented in the perisylvian, “higher-order” brain regions, including the inferior frontal, superior temporal, and parietal regions (Regev et al., 2013).

In contrast, other studies have reported that selective attention can improve the comprehension of sentences in the attended modality and induce changes in cortical activity patterns (Moisala et al., 2015; Regev et al., 2019; Wang and He, 2014). Regev et al. (2019) showed that selective attention can enhance the linguistic information flow of the attended modality, from early sensory areas along the processing hierarchy, converging in the perisylvian, “higher-order” brain regions.

Although the processing hierarchy of linguistic information has been suggested, in terms of both modality invariance and attention selectivity, independently, how these two “higher-order” areas are related to each other is yet to be determined (Figure 1A). The first hypothesis is that these areas overlap, forming a unified area that represents modality-invariant and attention-selective information (Figure 1A, left). The second hypothesis is that functionally distinct areas operate independently (Figure 1A, right). The third hypothesis exists between these two extremes: some areas represent both modality invariance and attention selectivity, whereas other areas exclusively represent one or the other (Figure 1A, center). Which types of linguistic information (semantic or phonemic) contribute to these properties also remains unknown.

**Figure 1.**
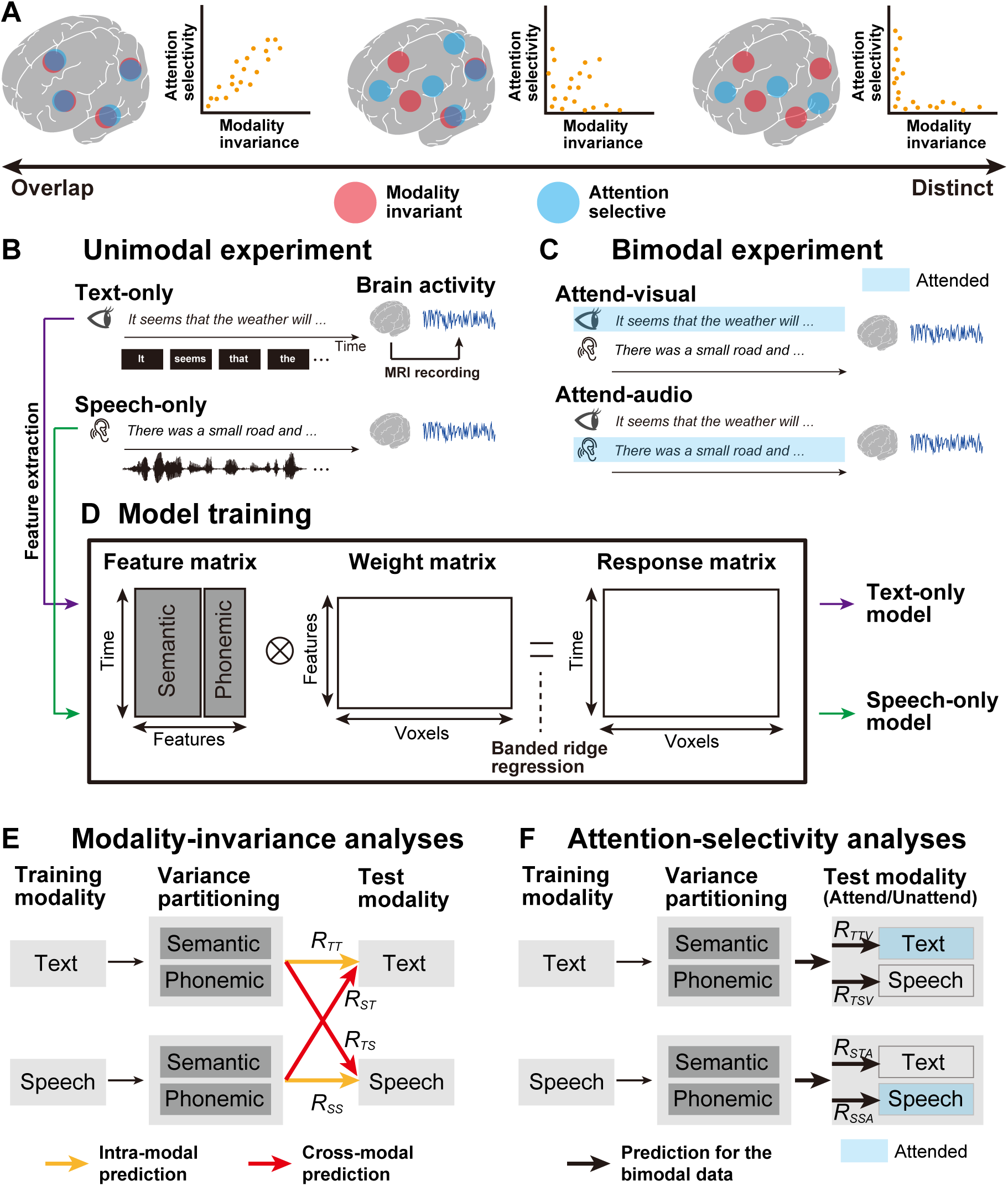
Schematic image of the experimental paradigm and the encoding modeling. (**A**) Three possible hypotheses are possible regarding the overlap between brain regions associated with modality-invariant linguistic representations (red) and those modulated by selective attention (blue). Brain regions showing these higher-order properties could be overlapping (left), independent (right), or partially overlapping (center). (**B**) Experimental design. During the unimodal experiment (left panel), participants passively listened to auditory stimuli, in the Speech-only condition, or read a written text, in the Text-only condition. Brain activity was measured using functional magnetic resonance imaging (fMRI). The original Japanese stimuli were translated into English for the purposes of intelligibility. (**C**) During the bimodal experiment (right panel), visual and auditory stimuli were presented to participants, simultaneously. The participants selectively attended to the visual (Attend-visual condition) or auditory (Attend-audio condition) modality. The stimuli in the attended modality are highlighted in blue. (**D**) Semantic and phonemic features were extracted from the text and speech stimuli used during the unimodal experiment, and encoding models were separately trained, using the brain activity of the training dataset (text-only and speech-only models). For model training, using a concatenated matrix of semantic and phonemic features, we used a banded ridge regression (see Methods). (**E**) For modality invariance analyses, trained unimodal models were used to predict brain activity in the test dataset from the unimodal experiments, in both intra-modal (yellow arrows) and cross-modal (red arrows) manners. The prediction accuracy notations are described with each arrow (see Table 1 for a description of all notations). Semantic and phonemic components were separated using variance partitioning analysis (see Methods). (**F**) For attention selectivity analyses, the trained unimodal models were used to predict brain activity during the bimodal experiment, and semantic and phonemic features were extracted from both attended and unattended modalities. Stimuli in the attended modality are highlighted in blue.

To address these issues, 7-hour, functional magnetic resonance imaging (fMRI) experiments were conducted, during which unimodal and bimodal language stimuli were presented. During the unimodal experiments (Figure 1B), six participants either listened to spoken narratives (Speech-only condition) or read transcribed narratives (Text-only condition). Meanwhile, during the bimodal experiment (Figure 1C), both speech and text were presented simultaneously, and the participants were asked to attend to either the speech or the text (Attend-audio or Attend-visual conditions, respectively). Data from the unimodal experiment were used to evaluate modality invariance, whereas the data from the bimodal experiment were used to evaluate attention selectivity.

**Table 1.** Notation of prediction accuracy for all combinations of conditions, for both the training and test datasets

| Modality invariance analyses (unimodal experiments) |  |  |  |  |
| --- | --- | --- | --- | --- |
| Training condition | Test condition and feature modality |  |  |  |
|  | Text-only condition<br>(Features from the text) |  | Speech-only condition<br>(Features from speech) |  |
| Text-only condition | $R_{TT}$ | | $R_{TS}$ | |
| Speech-only condition | $R_{ST}$ | | $R_{SS}$ | |
| Attention selectivity analyses (bimodal experiments) |  |  |  |  |
|  | Attend-visual condition |  | Attend-audio condition |  |
|  | Features from the text (attended) | Features from speech (unattended) | Features from the text (unattended) | Features from speech (attended) |
| Text-only condition | $R_{TTV}$ | $R_{TSV}$ | | |
| Speech-only condition | | | $R_{STA}$ | $R_{SSA}$ |

In order to evaluate the brain representations of multiple linguistic features, quantitatively, we used voxel-wise encoding models (Naselaris et al., 2011) (Figure 1D). By using this approach, brain activity can be modeled by a combination of features that are extracted from the presented stimuli. Researchers have adopted this approach to comprehensively examine semantic representations (de Heer et al., 2017; Deniz et al., 2019; Huth et al., 2016; Van Uden et al., 2018; Zhang et al., 2020), visual object category representations (Çukur et al., 2016; Huth et al., 2012), and how attention modulates representations (Çukur et al., 2013). Using this modeling approach, under cross-modal and multiple-attention conditions, we elucidated a quantitative relationship between modality-invariant representations (Figure 1E) and attentional modulation (Figure 1F), in a feature-specific manner.

## Materials and Methods

### Participants

Six healthy participants (referred to as ID01–ID06; ages 22–29; all native Japanese; two females), with normal vision and hearing, participated in the fMRI experiments. Participants were all right-handed, as measured using the Edinburgh inventory (Oldfield, 1971) (laterality quotients, 62.5– 100). Informed consent was obtained from all participants, prior to their participation in the study. This experiment has received approval from the Ethics and Safety Committee of the National Institute of Information and Communications Technology, in Osaka, Japan.

### Stimuli and tasks

We selected 20 narrative stories from the Corpus of Spontaneous Japanese (Maekawa, 2003), of which, 14 narratives were used during the training runs, for both Text-only and Speech-only conditions. One narrative was only used in the test runs of the Text-only condition, one narrative was only used in the test runs of the Speech-only condition, two narratives were only used in the test run of the Attend-visual condition, and two narratives were only used in the test run of the Attend-audio condition. All test runs were conducted twice. We used different narratives during each test runs, in order to avoid adaptations to the redundant presentation of the same content.

All narratives were originally recorded in the auditory modality. Sound signals were controlled by their root mean square and were only used in the Speech-only, Attend-visual, and Attend-audio conditions. Visual stimuli used for the Text-only, Attend-visual, and Attend-audio conditions were generated by presenting each spoken segment on the center of the screen. The onset of each visual segment has matched the onset of the corresponding segment in the spoken narrative. The average duration of the spoken narratives [mean ± standard deviation (SD)] was 673 ± 70 s.

During the Speech-only condition, participants were asked to fixate on a fixation cross-presented on the center of the screen and listened to spoken narratives, through MRI-compatible ear tips. Meanwhile, during the Text-only condition, participants read the transcribed narratives, which were displayed on the center of the screen, using a rapid, serial, visual presentation method. During the Attend-audio condition, participants listened to the spoken narratives, through MRI-compatible ear tips, and were instructed to ignore the text that was displayed simultaneously. Participants were asked not to close their eyes and were further instructed to fixate on the center of the screen. During the Attend-visual condition, participants were instructed to read the transcribed narratives displayed on the center of the screen, while ignoring the simultaneously presented spoken narratives.

At the beginning of each run, 10 s of dummy scans were acquired, during which the fixation cross was displayed, and these dummy scans were later omitted from the final analysis to reduce noise. We also obtained 10 s of scans at the end of each run, during which the fixation cross was displayed, and these were included in the analyses. In total, 36 fMRI runs were performed for each participant. Among these, 28 runs were used for model training (14 each, under the Speech-only and Text-only conditions), whereas 8 runs were performed for model testing (2 each, under the Text-only, Speech-only, Attend-visual, and Attend-audio conditions). For each participant, the experiments were executed over the course of 7 days, with 4–6 runs performed each day.

Participants were informed prior to the fMRI scan that, after each run, they would be asked to answer 10 questions relevant to the stimulus on which they were instructed to concentrate (the attended stimulus). However, the actual questionnaire that was administered after the fMRI scans included 10 questions that were relevant to both the attended and unattended stimuli. This surprise was intended so that participants would concentrate on understanding the instructed modality while ignoring the distractive one.

### MRI data acquisition

This experiment was conducted on a 3.0 T MRI scanner (MAGNETON Prisma; Siemens, Erlangen, Germany), with a 64-channel head coil. We scanned 72 2.0-mm-thick interleaved, axial slices, without a gap, using a T2-weighted, gradient-echo, multiband, echo-planar imaging (MB-EPI) sequence (Moeller et al., 2010) [repetition time (TR) = 1,000 ms, echo time (TE) = 30 ms, flip angle (FA) = 62°, field of view (FOV) = 192 × 192 mm^2^, voxel size = 2 × 2 × 2 mm^3^, multiband factor = 6]. The number of volumes collected was determined to be different for each run, depending on the stimuli length, of (mean ± SD) 693 ± 70 s (including the 10 s of initial dummy scans and the 10 s of fixation scans at the end of each run). For anatomical reference, high-resolution T1-weighted images of the whole brain were also acquired from all participants, using a magnetization-prepared rapid acquisition gradient-echo sequence (MPRAGE, TR = 2,530 ms, TE = 3.26 ms, FA = 9°, FOV = 256 × 256 mm^2^, voxel size = 1 × 1 × 1 mm^3^).

### Semantic features

To quantitatively evaluate the brain representations of the presented semantic information, in a data-driven manner, we extracted the semantic features from each narrative stimulus, using Wikipedia2Vec (Yamada et al., 2018) (https://wikipedia2vec.github.io/wikipedia2vec/). Wikipedia2Vec has been used to embed words and entities into distributed representations, based on the skip-gram model (Mikolov et al., 2013). All transcribed narrative segments were further segmented into words and were morphologically parsed, using MeCab (https://taku910.github.io/mecab/). Individual word segments were projected into the 300-dimensional space (i.e., word vectors with 300 elements) and were later assigned to the mean time point between the onset and offset of target segments, with 40 Hz. Time points without any word vector assignments were defined as 0. The resultant concatenated vectors were downsampled to 1 Hz.

### Phonemic features

To compare the predictability of brain activity, based on semantic features, with that of other non-semantic linguistic features, we also extracted phonemic features from each narrative stimulus. By using the Julius speech recognition software (Lee et al., 2001), an onset of each phoneme included in each spoken narrative was extracted. Each phoneme was then temporally aligned, based on the estimated onset. In total, 39 phonemes were extracted using the phoneme alignment procedure, which was transformed into one-hot vectors. Each one-hot vector was assigned to the mean time point between the onset and the offset of the target phoneme, with 40 Hz. Time points without any word vector assignments were defined as 0. The resulting concatenated vectors were downsampled to 1 Hz.

### fMRI data preprocessing

Motion corrections for each run were performed using the Statistical Parametric Mapping Toolbox (SPM8). All volumes were aligned using the first echo-planar imaging (EPI) result for each participant. Low-frequency drift was removed, using a median filter, with a 120-s window. The response for each voxel was then normalized, by subtracting the mean response and scaling the response to the unit variance. We used FreeSurfer (Dale et al., 1999; Fischl et al., 1999) to identify cortical surfaces, based on anatomical data, and to register them against voxels for functional data. For each participant, voxels identified throughout the whole brain were used in the analysis.

### Encoding model fitting

The cortical activities measured in each voxel were fitted with a set of linear temporal filters, which captured the slow hemodynamic responses, which were coupled with brain activity (Nishimoto et al., 2011). In order to examine how the different linguistic features were associated with cortical activity patterns, we modeled brain activity using two linguistic features (phonemic and semantic). A semantic feature matrix, F_1_ [T × 6N_1_], was modeled by concatenating sets of [T × N_1_] semantic feature matrices, with six temporal delays of 2 to 7 s each (T = # of samples; N_1_ = # of features). Similarly, the phonemic feature matrix, F_2_ [T × 6N_2_], was modeled using concatenating sets of [T × N_2_], using phonemic feature matrices, with six temporal delays. A cortical response, X [T × V], was then modeled by the concatenated feature matrices of F_1_ and F_2_, multiplied by the concatenated weight matrices, W_1_ [6N_1_ × V] and W_2_ [6N_2_ × V] (V = # of voxels):

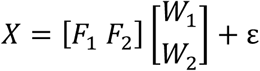

where ε is isotropic gaussian noise. In analyzing the predictive performance of the two linguistic models exclusively, we used banded ridge regression on the training dataset to obtain the weight matrices, W_1_ and W_2_ (Nunez-Elizalde et al., 2019). Specifically, weight matrices were estimated by solving the following equation, with regularization parameters α_1_ and α_2_:

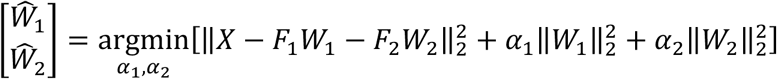

The solution to this equation is:

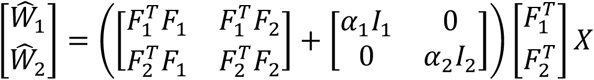

where I_1_ and I_2_ are identity matrices of the sizes [6N_1_ × 6N_1_] and [6N_2_ × 6N_2_], respectively. The training dataset consisted of 9,815 samples, under both Speech-only and Text-only conditions. An optimal regularization parameter was assessed in each voxel using 10-fold cross-validation.

The test dataset consisted of 619 Speech-only samples, 617 Text-only samples, 613 Attend-audio samples, and 623 Attend-visual samples. Differences in sample sizes in the test dataset can be attributed to the various durations of the naturalistic narrative story stimuli. Two repetitions of the test dataset were averaged to increase the signal-to-noise ratio. Prediction accuracy was calculated, using Pearson’s correlation coefficient, between the predicted signal and the measured signal in the test dataset. Significant differences were computed using a one-sided Student’s *t*-test. The resulting *p*-values were corrected for multiple comparisons within each participant, using the false discovery rate (FDR) procedure (Benjamini and Hochberg, 1995).

### Variance partitioning analysis

To assess the predictive performances of semantic and phonemic features separately, we performed a variance partitioning analysis (de Heer et al., 2017; Lescroart et al., 2015). Predicted signals were estimated for each of the two separate models and the concatenated model, as follows:

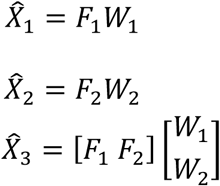

Explained variances (coefficients of determination) were also estimated for each of the two separate models and the concatenated model, as follows:

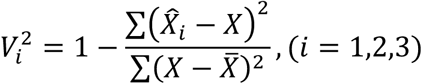

where 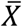 represents a mean response in the test dataset (across time). Prediction accuracies for every single model (*R*_*1*_ and *R*_*2*_, for the semantic and phonemic features, respectively) were obtained by subtracting the coefficient of determinant calculated for a targeted single model from that calculated for the concatenated model, as follows:

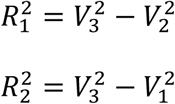

To make the predicted performance comparable with those reported by previous studies (de Heer et al., 2017; Deniz et al., 2019; Huth et al., 2016, 2012), the square root of the explained variances was calculated. To obtain a null distribution of the explained variances, we calculated *R*_*1*_ and *R*_*2*_ values for all cortical voxels, based on the originally predicted responses and a random permutation of the actual responses (a permutation of the temporal order) from the test dataset. The resulting *p*-values (one-sided) were corrected for multiple comparisons using the FDR procedure (Benjamini and Hochberg, 1995).

### Modality invariance

To quantify how the unimodal models explained brain activity in each voxel, regardless of the presentation modality, we defined an *MI* value, which consisted of two components: *D*_*T*_ and *D*_*S*_. *D*_*T*_ has been defined as a degree of predictability for the Text-only test dataset, regardless of the training modality:

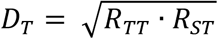

where *R*_*TT*_ and *R*_*ST*_ are the intra-modal prediction accuracy for the text-only model and the cross-modal prediction accuracy calculated for the speech-only model when applied to the test dataset for the Text-only condition, respectively (see Table 1 for all notations of prediction accuracies). Note that the R_**_ values correspond to the *R*_*1*_ or *R*_*2*_ values described in the previous subsection, depending on the linguistic features used (i.e., semantic or phonemic). Similarly, *D*_*S*_ is defined as the degree of predictability calculated for the Speech-only test dataset, regardless of the training modality:

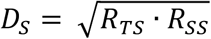

where *R*_*SS*_ and *R*_*TS*_ are the intra-modal prediction accuracy by the speech-only model and the cross-modal prediction accuracy calculated for the text-only model when applied to the test dataset for the Speech-only condition, respectively. For all voxels showing negative prediction accuracies, the prediction accuracy was set to 0 to avoid obtaining imaginary values. Modality invariance (*MI*) was then calculated for each voxel as a geometric mean between *D*_*S*_ and *D*_*T*_, as follows:

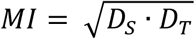

A high *MI* value indicated that the target linguistic features are represented in a modality-independent manner. The significance of each MI value was assessed using a permutation test (one-sided), corrected for multiple comparisons using the FDR procedure (Benjamini and Hochberg, 1995).

### Modality specificity

To quantify how the unimodal models explained brain activity that was specific for a single modality, we defined modality specificity, which was calculated in each voxel for each modality (MS_T_ for the Text-only condition and MS_S_ for the Speech-only condition) as the difference between the intra-modal and cross-modal prediction accuracies:

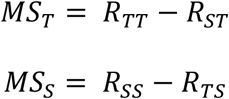

A high *MS* value indicated that the target linguistic features are represented specifically according to the target modality. Significance and FDR corrections for multiple comparisons were calculated as described for the *MI* values.

### Attention selectivity

To quantify how each cortical voxel was affected by selective attention, we defined an *AS* value, consisting of two components, *A*_*V*_, and *A*_*A*_, which indicated the augmentation of prediction accuracies according to the application of selective attention to the visual and auditory modalities, respectively. To calculate *A*_*V*_, the text-only model was tested on the test dataset acquired under the Attend-visual condition. The prediction accuracies contrasted the features from the visual (attended) and the auditory (unattended) modalities, as follows:

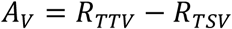

where *R*_*TTV*_ and *R*_*TSV*_ represent the prediction accuracies calculated based on the features from the visual (attended) and auditory (unattended) modalities, respectively (see Table 1 for all notations of prediction accuracies). Similarly, to calculate *A*_*A*_, the speech-only model was tested on the test dataset acquired under the Attend-audio condition. The prediction accuracies contrasted the features from the auditory (attended) and visual (unattended) modalities, as follows:

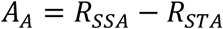

where *R*_*SSA*_ and *R*_*STA*_ are the prediction accuracies calculated based on the features from the auditory (attended) and visual (unattended) modalities, respectively. Voxels showing negative *A*_*V*_ and *A*_*A*_ values were set to 0. Attention selectivity (*AS*) was calculated as the geometric mean of *A*_*V*_ and *A*_*A*_, as follows:

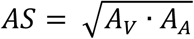

*AS* is high when the features extracted from the attended modality always predict brain activity better than those of the unattended modality. For comparison purposes, voxels that showed significant prediction accuracies in the unimodal models (using the minimum statistic as a threshold) were also used to visualize the *AS* results. The calculation of statistical significance and FDR corrections for multiple comparisons were performed as described for the *MI* values.

### Anatomical ROI analysis of modality invariance and attention selectivity

To quantify how different cortical regions display modality invariance and attention selectivity, we calculated ratios between voxels with exclusively positive MI values (*MI-only voxels*), voxels with exclusively positive AS values (*AS-only voxels*), or voxels showing both positive MI and AS values (*shared voxels*) and voxels showing either positive MI or AS values across 148 anatomical regions of interest (ROIs), based on the Destrieux cortical atlas (Destrieux et al., 2010). To focus on the cortical regions associated with linguistic information, we selected ROIs that contained a relatively large number of voxels with positive MI or AS values (> 20 % and > 50 voxels within the target ROI).

## Results

### The semantic encoding model predicted brain activity, regardless of modality

To confirm that the encoding models successfully captured brain activity during the unimodal experiments (Figure 1B), we performed a series of intra-modal encoding modeling tests and examined the modeling accuracy using a test dataset from the same modality as the training dataset (Figure 1E, yellow). To quantifiably evaluate the predictability of brain activity, based on different linguistic information, we extracted both semantic and phonemic features from the narrative stimuli. We exclusively evaluated the effects of either semantic or phonemic features by combining banded ridge regression (Nunez-Elizalde et al., 2019) with variance partitioning analysis (de Heer et al., 2017; Lescroart et al., 2015) (see Methods for details).

We first trained encoding models using the data from the Text-only condition (text-only model) and predicted brain activity using the text-only test dataset. Using semantic features, we found that the text-only model significantly predicted activity in the perisylvian regions, including the superior temporal, inferior frontal, and inferior parietal cortices (blue or white in Figure 2A). Similarly, we trained encoding models using the data from the Speech-only condition (speech-only model) and predicted brain activity using the speech-only test dataset (blue or white in Figure 2B). The speech-only model also significantly predicted activity in the perisylvian regions.

**Figure 2.**
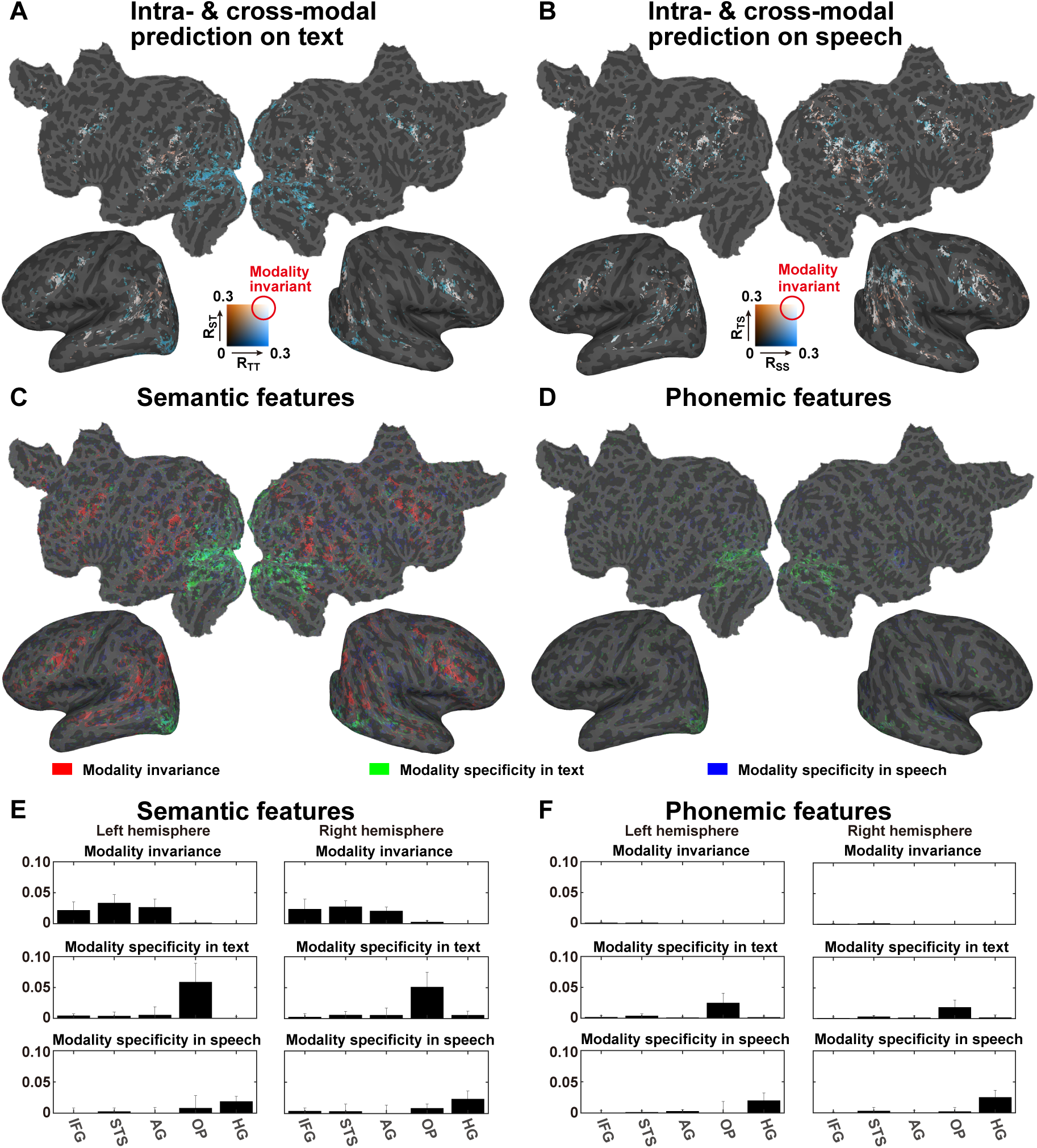
Effects of stimulus modality. (**A**) Comparison of the prediction accuracies of the unimodal models when applied to a Text-only test dataset using semantic features. The intra-modal prediction accuracies of the Text-only model (denoted as *R*_*TT*_, blue) and the cross-modal speech-only model (denoted as *R*_*ST*_, orange) are mapped onto the cortical surface of participant ID01. Regions in which activity was predicted, regardless of the stimulus modality, are shown in white. Only significant regions (*p* < 0.05 with FDR correction) are shown. (**B**) Comparison between the prediction accuracies of the unimodal models for a Speech-only test dataset, using semantic features. Intra-modal prediction accuracy, using the speech-only model (denoted as *R*_*SS*_, blue), and the cross-modal prediction accuracy, using the text-only model (denoted as *R*_*TS*_, orange), are mapped onto the cortical surface. (**C** and **D**) Modality invariance (MI, red), modality specificity for text (MS_T_, green), and modality specificity for speech (MS_S_, blue) were calculated by taking the geometric mean of the intra-modal and cross-modal prediction accuracies, using semantic features (**C**) or phonemic features (**D**). Significant regions (*p* < 0.05 with FDR correction) were shown in each color (see Figures S1 and S2 for the other participants). (**E** and **F**) The mean MI, MS_T_, and MS_S_ values were extracted from the following anatomical regions of interest: inferior frontal gyrus (IFG), superior temporal sulcus (STS), angular gyrus (AG), occipital pole (OP), and Heschl’s gyrus (HG), averaged across six participants, for both the left and right hemispheres, using semantic features (**E**) or phonemic features (**F**). Error bar, SD.

We next examined whether the unimodal models captured modality-invariant representations by performing cross-modal encoding modeling, during which we examined the modeling accuracy using a test dataset obtained from a different modality than the training dataset (Figure 1E, red). A speech-only model was used to predict the brain activity with a text-only test dataset (*p* < 0.05 with FDR correction, red or white in Figure 2A), which showed significant prediction accuracy in the perisylvian regions. Similarly, a text-only model was used to predict brain activity with a speech-only test dataset, which also displayed significant prediction accuracy in the perisylvian regions (red or white in Figure 2B). The overlap between the intra-modal and cross-modal prediction performances displayed a clear contrast between the cortical organization associated with modality-specific representation in the early sensory regions (i.e., the superior temporal and occipital regions) and that associated with the modality-invariant representation in the perisylvian regions.

To identify those regions that activate during modality-invariant representations of linguistic information, we further calculated the modality invariance (MI) value, by combining the intra-modal and cross-modal prediction accuracies, using semantic features. The MI values were determined to be significantly larger than 0 in the perisylvian regions (*p* < 0.05 with FDR correction, red in Figure 2C), indicating that these regions are associated with modality-independent semantic representations. To identify those regions associated with modality-specific representations, we calculated the modality specificity of text (MS_T_) and speech (MS_S_). Even though significantly larger MS_T_ values could be observed in the visual cortex (*p* < 0.05 with FDR correction, green in Figure 2C), the MS_S_ values were found to significantly increase in the auditory cortex (*p* < 0.05 with FDR correction, blue in Figure 2C). Phonemic features were also associated with significant MS_T_ values in the visual cortex and with MS_S_ values in the auditory cortex. In contrast with semantic features, however, phonemic features were associated with very small modality invariance values across the cortex (Figure 2D), indicating that modality-invariant representations of linguistic information are predominantly conveyed by semantic features.

To evaluate the modality invariance associated with each cortical region, we extracted modality invariance values for five anatomical regions of interest (ROIs), averaged across all six participants (Figures 2E and F). For the anatomical ROIs, we selected three perisylvian regions: the inferior frontal gyrus (IFG), the superior temporal sulcus (STS), and the angular gyrus (AG). Activity in these regions has frequently been reported in previous neuroimaging studies examining language (Hagoort, 2019; Price, 2010; Skeide and Friederici, 2016). We also selected two sensory ROIs, at the occipital pole (OP) and Heschl’s gyrus (HG), which process early sensory components in the visual and auditory modalities, respectively. We found that modality invariance values for semantic features were larger in the three perisylvian regions than those in the sensory regions (Cohen’s *d* = 1.45, which was calculated between the average of the three perisylvian regions and the average of the two sensory regions). In contrast, the MS_T_ values for semantic features in the three perisylvian regions were found to be smaller than that in the OP (*d* = 1.73), and the MS_S_ values for semantic features in the three perisylvian regions were smaller than that in the HG (*d* = 2.67). These results also indicated that only semantic features are represented in the perisylvian regions in a modality-invariant manner, whereas modality-specific information for both the visual and auditory domains are represented in the primary sensory areas, regardless of linguistic features.

### Effect of selective attention on model prediction performance

To examine whether selective attention affects the cortical representations of linguistic information, we conducted bimodal experiments, during which speech and text were simultaneously presented and participants were asked to selectively attend to only one of the two modalities (Figure 1C). During the Attend-visual condition, we extracted semantic features from both the attended (visual) and unattended (auditory) modalities, which were presented simultaneously. The prediction accuracies were calculated by applying a text-only model, with features in each of the attended and unattended modalities (Figure 1F, top). We found increased prediction accuracy across the cerebral cortex when using semantic features from the attended modality (Figure 3A, orange), compared with those from the unattended modality (blue). Similarly, a speech-only model was tested using the data collected during the Attend-audio condition. We again found larger prediction accuracy across the cerebral cortex when using semantic features from the attended modality (Figure 3B, orange), compared with those from the unattended modality (blue). Cross-modal prediction accuracies were not calculated during this procedure, and we evaluated modality invariance and attention selectivity separately.

**Figure 3.**
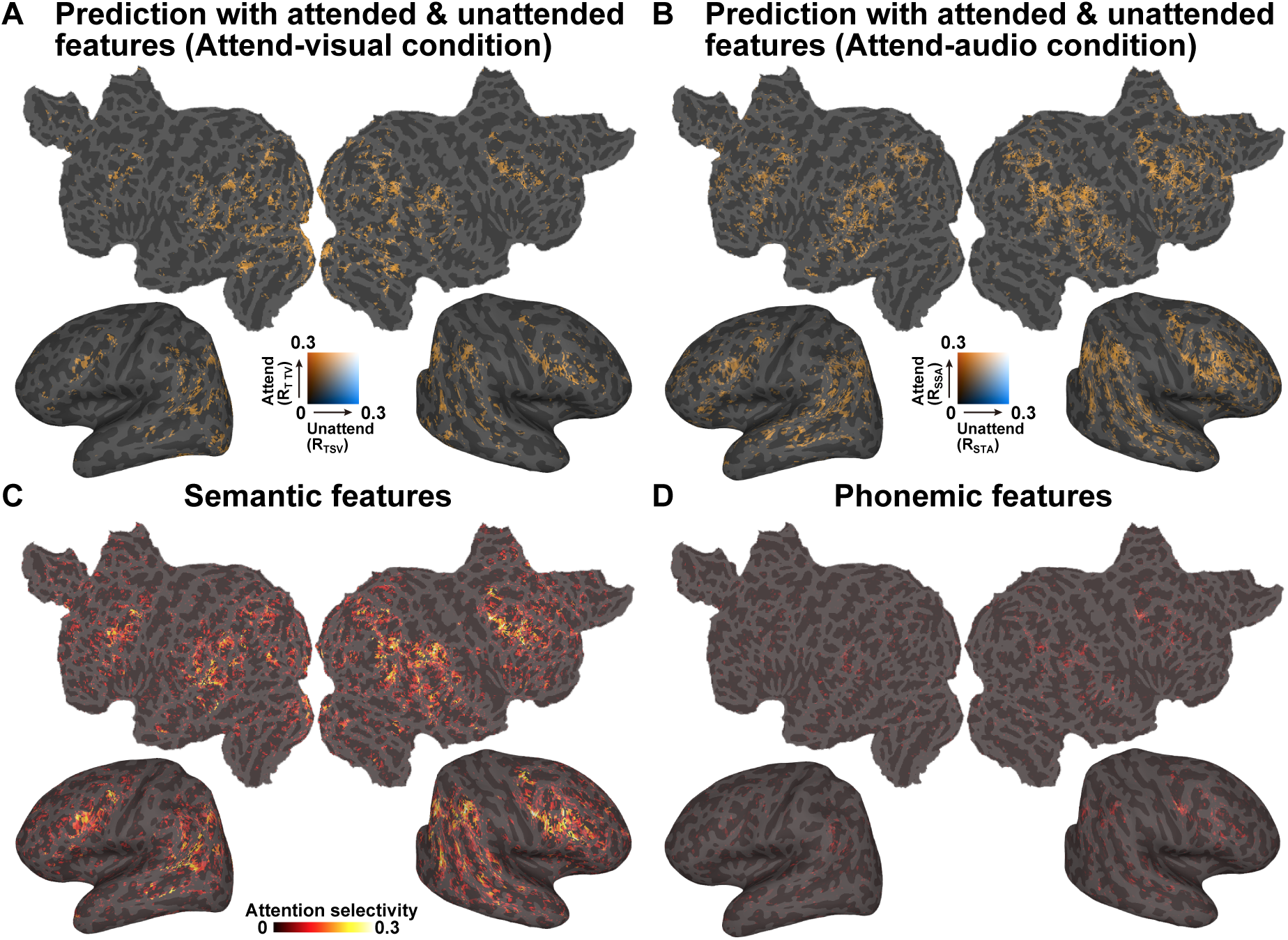
Effects of selective attention. (**A**) Prediction accuracies, using a text-only model on the Attend-visual condition dataset, mapped onto the cortical surface of participant ID01. Two prediction accuracies, associated with semantic features, were extracted from the attended modality (visual, orange) and the non-attended modality (audio, blue) and compared (*R*_*TTV*_ and *R*_*TSV*_, respectively). Only significant regions (*p* < 0.05 with FDR correction) are shown. (**B**) Comparison of the prediction accuracies using a speech-only model on the Attend-audio condition (*R*_*SSA*_ and *R*_*STA*_). **(C** and **D)** Attention selectivity was calculated by subtracting the prediction accuracy for the unattended condition from that for the attended condition and was mapped onto the cortical surface of participant ID01, using semantic features (**C**) or phonemic features (**D**) (see Figures S3 and S4 for the other participants).

To investigate which cortical regions were modulated by selective attention, we calculated attention selectivity (AS) by subtracting the prediction accuracy measured using unattended features from that calculated using attended features, within each modality (Figures 3A and B). Larger AS values were identified in the inferior frontal, middle temporal, and inferior parietal regions when using semantic features (Figure 3C). In contrast, we found that very small brain regions showed significant AS values when using phonemic features (Figure 3D).

### Select brain regions with modality-invariant semantic representations are affected by selective attention

An overlaid cortical map of the MI and AS values for semantic features (Figure 4A) indicated that some voxels specifically represented MI (red), whereas other voxels specifically represented AS (blue). A scatter plot of the cortical voxels clearly revealed three types of voxels associated with semantic features (Figure 4C for participant ID01; see Supplementary Figure S5 for the other participants), in which voxels associated with positive MI and 0 AS are colored in red (*MI-only voxels*; a mean ± SD voxel size across six participants of 5,016 ± 1,786), those associated with positive AS and 0 MI are colored in blue (*AS-only voxels*; 7,975 ± 2,121), and those associated with both positive MI and AS are colored in purple (*shared voxels*; 1,772 ± 1,153). Within the shared voxels, we found a positive correlation between AS and MI (Spearman’s correlation coefficients, *ρ* = 0.675 ± 0.035; Figure 4C). In contrast, when using phonemic features, relatively few shared voxels were found to have significant values (Figure 4D, 111 ± 48 voxels). However, we again found a positive correlation between AS and MI in these shared voxels (*ρ* = 0.670 ± 0.079).

**Figure 4.**
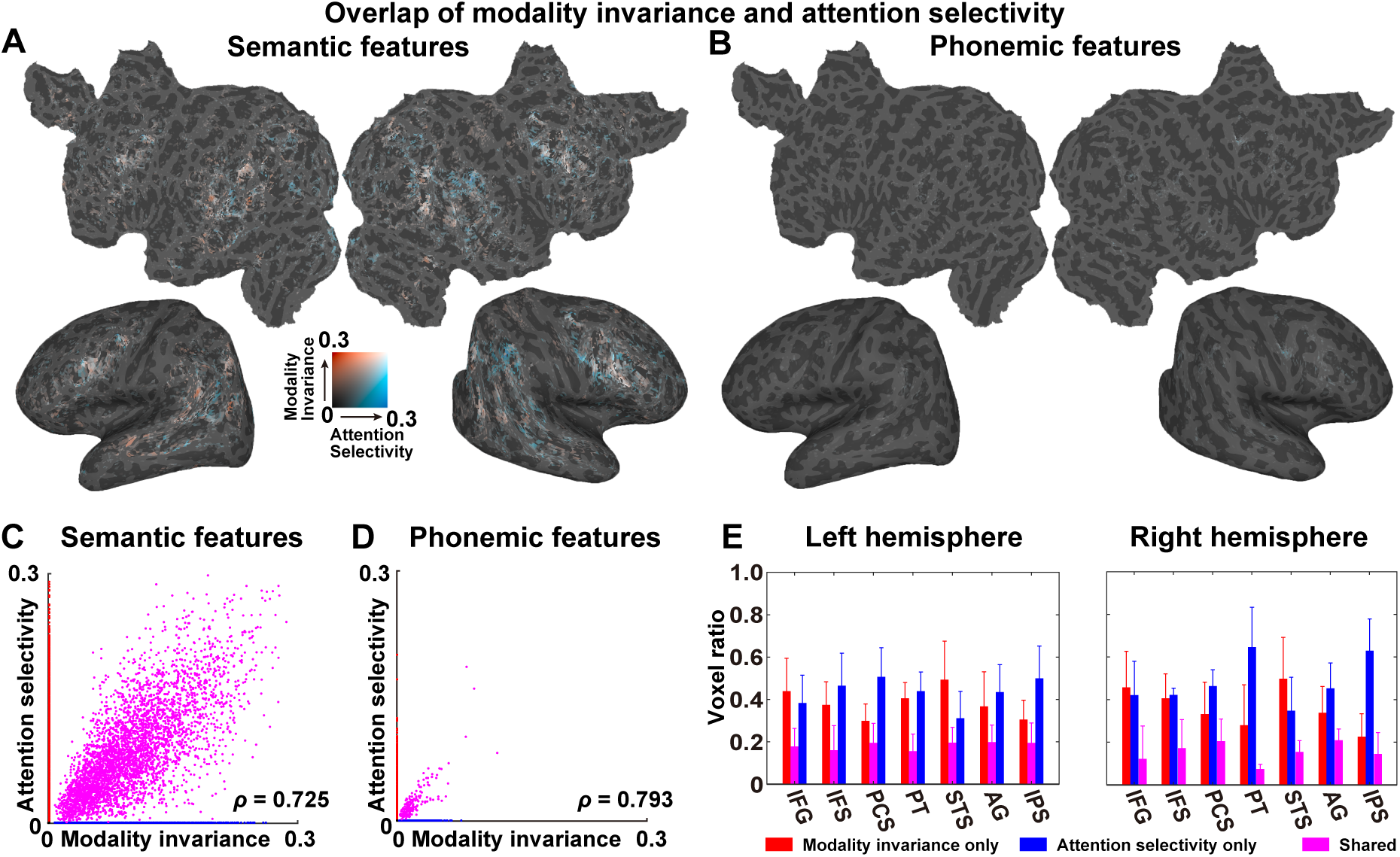
Partial overlap between the modality-invariant and attention-selective regions. **(A and B)** Modality invariance (MI, red) and attention selectivity (AS, blue) were mapped onto the cortical surface of participant ID01, using semantic features (**A**) and phonemic features (**B**). Only significant regions (*p* < 0.05 with FDR correction) are shown. Supplementary Figure S5 shows the plots for the other participants. (**C and D**) A scatter plot is shown for MI and AS, using semantic features (**C**) and phonemic features (**D**). Data were extracted from the cortical voxels of participant ID01. Data with positive MI and zero AS are colored in red. Data with positive AS and zero MI are colored in blue. Data with positive MI and AS are colored in purple. For each plot, a Spearman’s correlation coefficient (*ρ*) is displayed. **(E)** The ratios of voxels exclusively showing positive MI (red), those exclusively showing positive AS (blue), and those showing both MI and AS (purple) are plotted for seven anatomical regions of interest in the left (left panel) and right hemispheres (right panel). IFG, inferior frontal gyrus; IFG, inferior frontal sulcus; PCS, precentral sulcus; PT, planum temporale; STS, superior temporal sulcus; AS, angular gyrus; IPS, intraparietal sulcus. Error bar, SD.

To scrutinize the detailed cortical organization associated with MI and AS, we calculated the ratios between these three types of voxels and all voxels that display either positive MI or AS values across all of the anatomical ROIs when using semantic features (Figure 4E). Because both MI and AS were more densely associated with semantic features than with phonemic features (Figs. 4A–D), we focused on semantic features in this analysis. Seven bilateral perisylvian ROIs [IFG, inferior frontal sulcus (IFS), precentral sulcus (PCS), planum temporale (PT), STS, AG, and intraparietal sulcus (IPS)] contained a relatively large portion of voxels showing either significant MI or AS values (> 20 % and > 50 voxels within the target ROI). For all target ROIs, more MI-only and AS-only voxels appeared than shared voxels (*d* = 2.19 and *d* = 3.47, respectively). The ROIs showed different patterns, with more MI-only voxels found in the bilateral STS (left, *d* = 1.06; right, *d* = 0.77), whereas more AS-only voxels were found in the bilateral PCS (left, *d* = 1.68; right, *d* = 1.00), the right PT (*d* = 1.78) and the right IPS (*d* = 2.82).

## Discussion

In this current study, participants underwent fMRI experiments and were presented with either unimodal auditory or visual stimuli or with bimodal auditory and visual stimuli; they were later asked to selectively attend to only one modality. The unimodal model, using semantic features, was able to predict the activity in the bilateral inferior frontal, superior temporal, and inferior parietal regions, for both modalities. The involvement of these regions in language processing has been repeatedly suggested in many neuroimaging studies (Hagoort, 2019; Price, 2010; Skeide and Friederici, 2016). In contrast, the unimodal models using phonemic features were not able to predict modality-invariant activity. This result is consistent with the results of a previous study, by de Heer et al. (2017), which reported that encoding models using semantic features predicted larger brain regions than those predicted by phonemic models.

Differences in prediction accuracy, between attended and unattended features, can only be explained using the effects of attention as the only differences in the prediction were associated with these extracted features (attended vs. unattended). A behavioral questionnaire administered after each session confirmed that participants showed higher accuracy for understanding the semantic contents of the attended stimuli. This result suggested that semantic information is represented in the brain only when participants pay attention to the target stimuli, which was consistent with the finding that phonemic features produced significant AS values only in very small brain regions.

The brain regions that were well-predicted by cross-modal predictions and those regions that were modulated by selective attention demonstrated partial overlap; this agrees with the hypothesis depicted in Fig. 1A (center). Anatomical ROI analyses further showed that the overlapping regions were primarily located in perisylvian regions, such as the inferior frontal, superior temporal, and inferior parietal cortices. The modality invariance of semantic representations has been reported previously (Booth et al., 2002; Deniz et al., 2019); however, we show, for the first time, that this representation partially correlates with the effects of selective attention on perisylvian regions.

We also identified cortical voxels showing only modality invariance, as well as those showing only attention selectivity. These results suggested heterogeneity among cortical representations of semantic information. A recent study of the cocktail party effect (i.e., the simultaneous presentation of different auditory stimuli) showed that brain activity reflected the semantic information of attended words but not unattended words (Brodbeck et al., 2018). Other studies reported a reshaping of semantic representation of visual objects and behavioral categories by selective attention (Çukur et al., 2013; Nastase et al., 2017). Selective attention may affect some cortical semantic networks, in a modality-specific manner.

Modality-specific regions were primarily identified in the primary auditory cortex, for the auditory modality, and in the primary visual cortex, for the visual modality. These results are consistent with a previous study, which reported that higher-order brain regions were more strongly affected by selective attention than early sensory regions (Regev et al., 2019). Importantly, the modality-specificity values were found to be larger in the early sensory regions, for both semantic and phonemic features, whereas the modality-invariance values were larger in the perisylvian regions, only for semantic features. This model dependency may indicate that the processing of semantic information is more affected by selective attention, whereas phonemic features are primarily processed in early sensory regions and are not fully affected by selective attention.

Using an encoding model approach, we compared phonemic and semantic features for their predictabilities of brain activity, which provided detailed information that was not obvious in a previous study that reported increased inter-participant correlations among brain activities during selective attention (Regev et al., 2019). We found that encoding models associated with semantic features were more strongly affected by selective attention than the phoneme-based models, which is consistent with behavioral results showing that the understanding of semantic content was facilitated by selective attention.

Unimodal models using phonemic features have showed modality specificity, not only in the auditory cortex but also in the visual cortex. Although this finding may appear to contradict the idea of a “phoneme,” the MS_T_ values observed in the early visual cortex can be explained by phonemic orthography associated with the Japanese language. Because Japanese largely contains phonograms (i.e., *Hiragana* and *Katakana*), phonemic features may correlate with visually presented Japanese characters.

We used naturalistic, narrative stories and extracted linguistic information from both the auditory and visual stimuli. This approach can be applied to other linguistic features. Although we have only examined two linguistic models, which have both been used in previous studies (de Heer et al., 2017; Nishida and Nishimoto, 2018; Kivisaari et al., 2019), further applications examining different features, such as syntax, may further increase prediction accuracy and capture more profound representations across modalities. Further modeling approaches are necessary for the comprehensive evaluation of the cortical representation of linguistic information and the effects of selective attention.

## Acknowledgments

This work was supported by MEXT/JSPS KAKENHI [grant numbers JP20K07718, JP20H05023 in #4903 (Evolinguistics) for T.N., and JP15H05311 for S.N.], as well as JST CREST [JPMJCR18A5] and ERATO [JPMJER1801 (for S.N.)], for the partial financial support of this study. The funders had no role in the study design, data collection, and analysis, decision to publish, or preparation of the manuscript. T.N., H.Q.Y., and S.N. designed the study; T.N. and H.Q.Y. collected and analyzed the data; T.N., H.Q.Y., and S.N. wrote the manuscript. The authors declare no competing interests.

## Supplementary Information

### Behavioral results

To confirm that participants performed the selective attention task as instructed, we tested whether participants understood contents of the attended stimuli by using a post-experimental questionnaire. The average score of post-experimental questionnaire was 90.8% ± 4.9% (Mean ± SD) for the attended stimuli and 50.0% ± 4.5% for the non-attended stimuli (chance level = 50%), indicating that selective attention facilitated participants’ comprehension of semantic contents in the linguistic stimuli.

**Figure S1.**
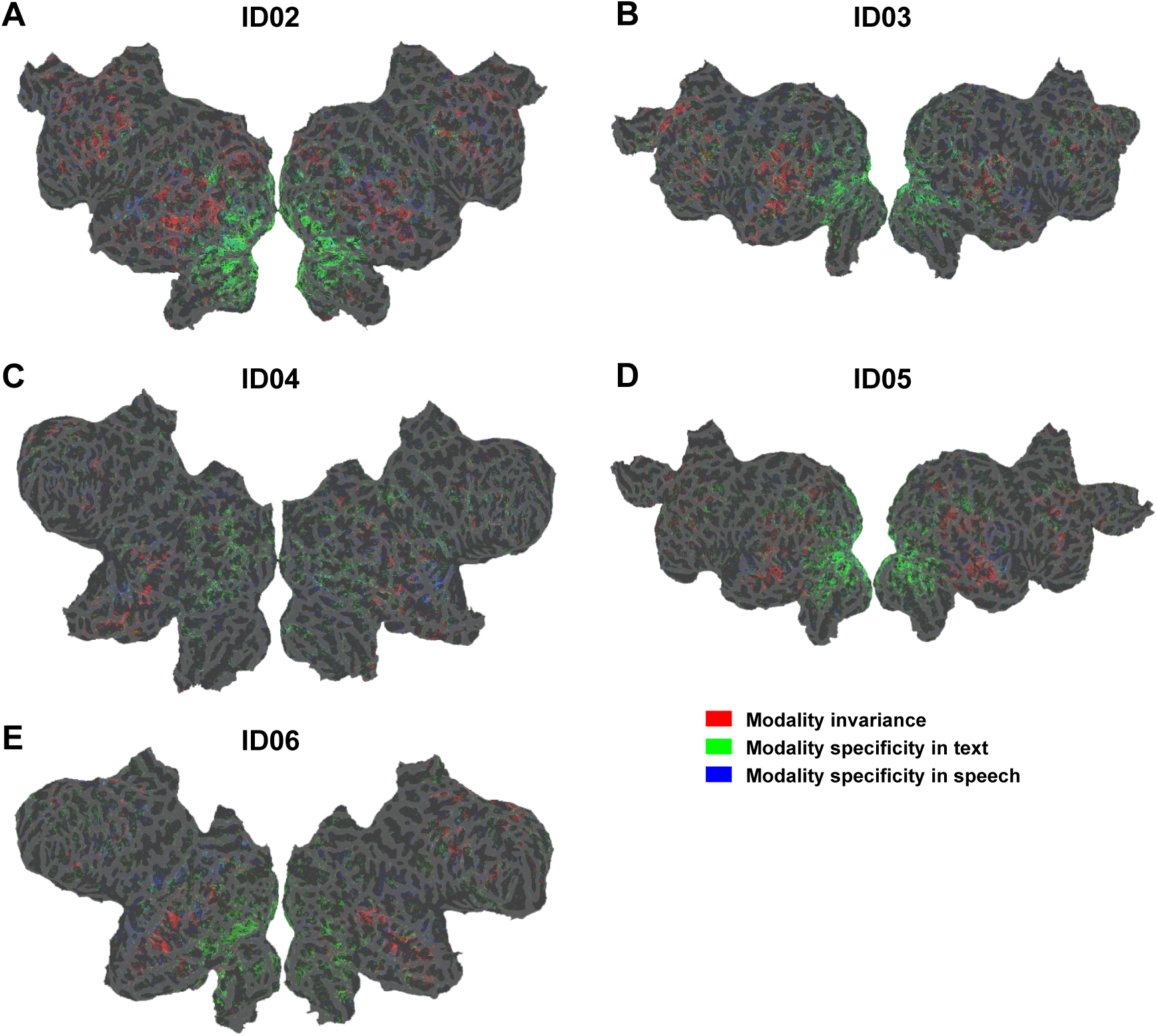
Modality invariance and modality specificity using semantic features. Modality-invariance (red), modality-specificity in text (green), and modality-specificity in speech (blue), calculated by taking the geometric mean of intra-modal and cross-modal prediction accuracies using semantic features, mapped onto the cortical surface of participant ID02-ID06. The significant regions (*p* < 0.05 with FDR correction) were shown in each color.

**Figure S2.**
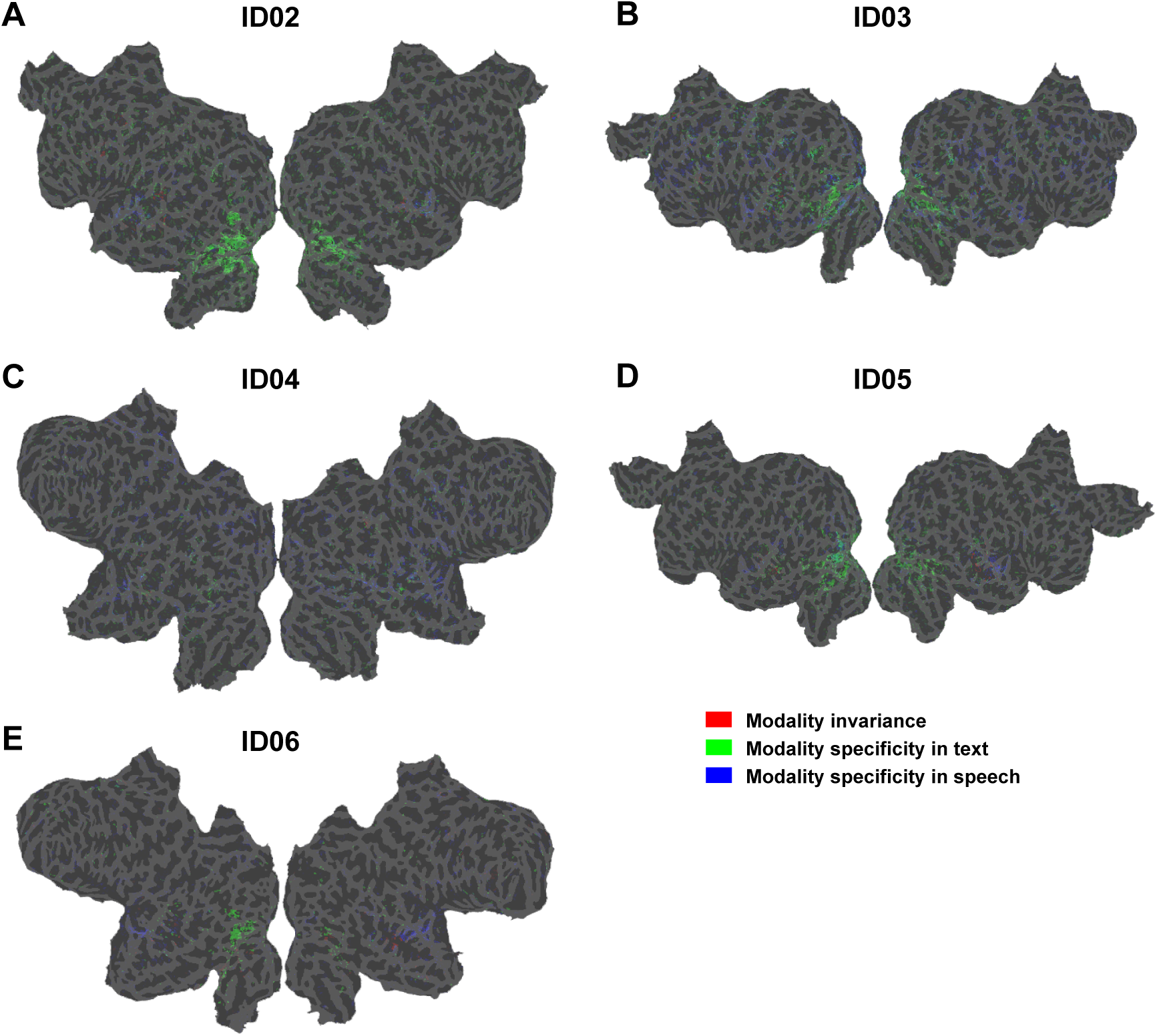
Modality invariance and modality specificity using phonemic features. Modality invariance (red), modality specificity in text (green), and modality specificity in speech (blue), calculated by taking the geometric mean of intra-modal and cross-modal prediction accuracies using phonemic features, mapped onto the cortical surface of participant ID02-ID06. Significant regions (*p* < 0.05 with FDR correction) were shown in each color.

**Figure S3.**
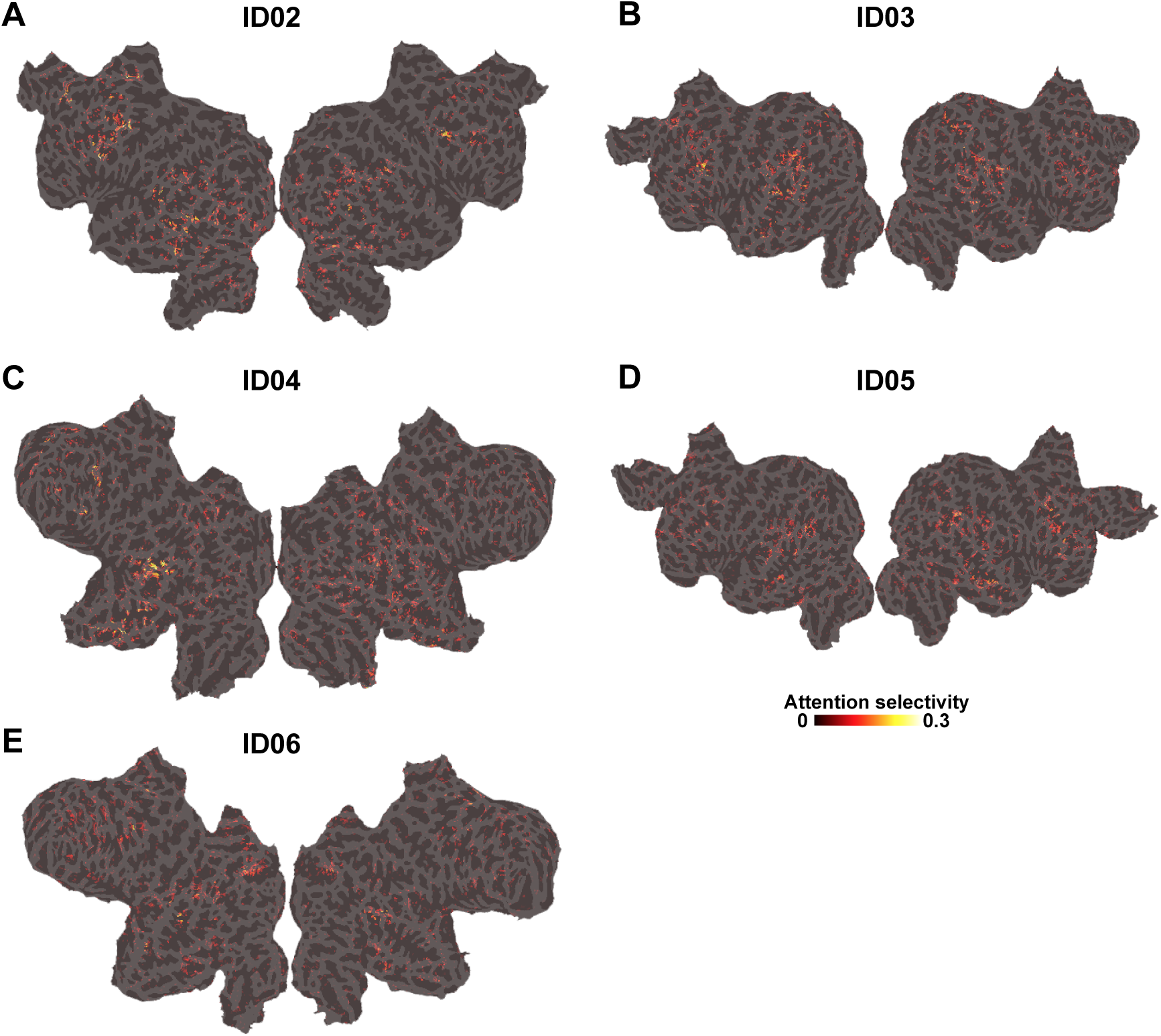
Attention selectivity using semantic features. Attention selectivity was calculated by subtracting prediction accuracy in unattended conditions from that in attended conditions, mapped onto the cortical surface of participant ID02-ID06 using semantic features. Only significant regions (*p* < 0.05 with FDR correction) were shown.

**Figure S4.**
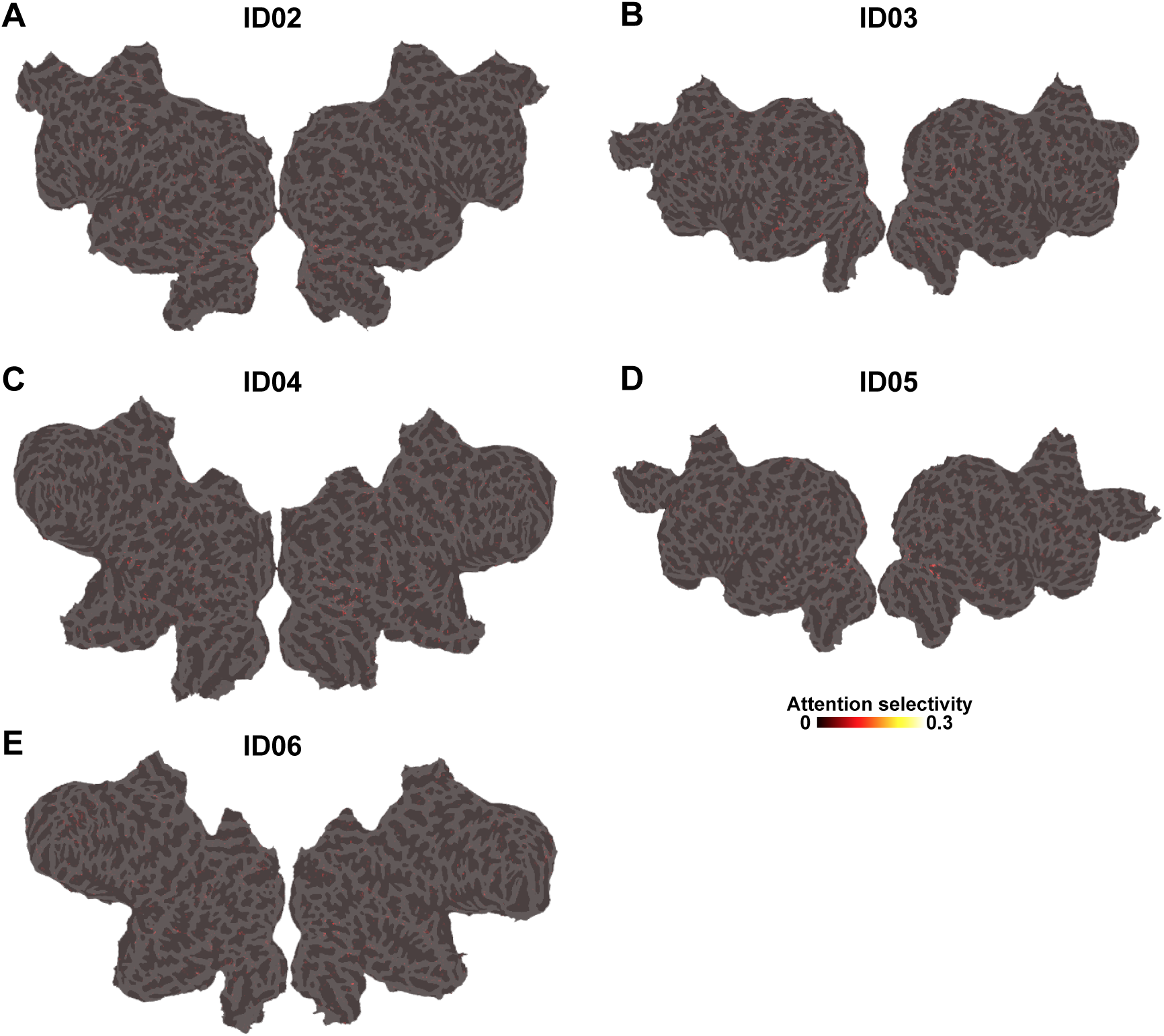
Attention selectivity using phonemic features. Attention selectivity was calculated by subtracting prediction accuracy in unattended conditions from that in attended conditions, mapped onto the cortical surface of participant ID02-ID06 using phonemic features. Only significant regions (*p* < 0.05 with FDR correction) were shown.

**Figure S5.**
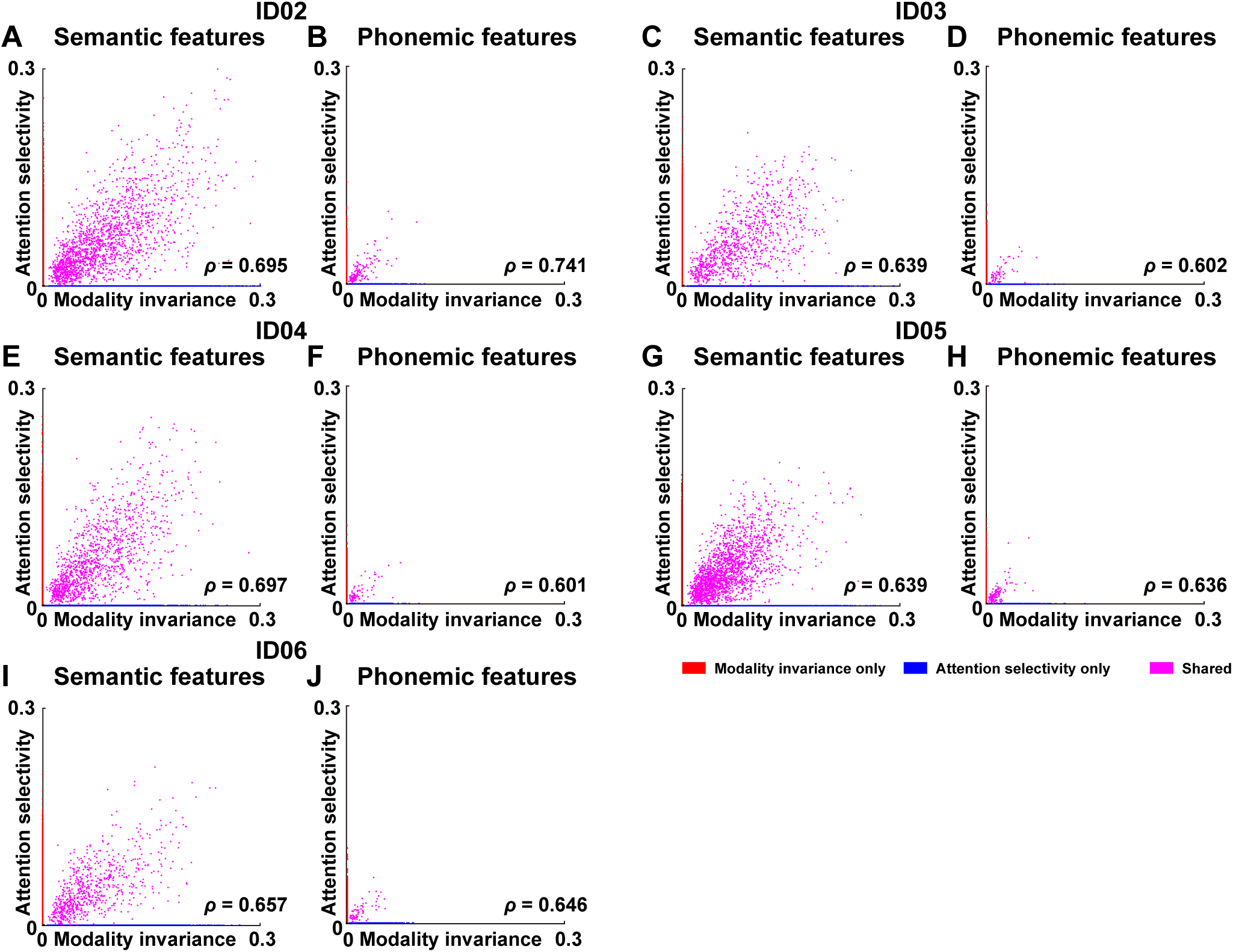
Scatter plots of modality-invariance and attention-selectivity. Scatter plot are shown for modality invariance and attention selectivity using semantic features and phonemic features, extracted from the cortical voxels of participant ID02-ID06. For each plot, a Spearman’s correlation coefficient (*ρ*) is displayed.

